# Profiling of chimeric RNAs in human retinal development with retinal organoids

**DOI:** 10.1101/2022.11.10.515982

**Authors:** Wen Wang, Xiao Zhang, Ning Zhao, Ze-Hua Xu, Kangxin Jin, Zi-Bing Jin

## Abstract

Chimeric RNAs have been found in both cancer and healthy human cells. They have regulatory effects on human stem/progenitor cell differentiation, stemness maintenance and central nervous system (CNS) development. However, their physiological functions in the retinal development remain unknown. Based on the human embryonic stem cells (hESC)-derived retinal organoids (ROs) spanning from day 0 to day 120, we present the expression atlas of chimeric RNAs throughout the developing ROs. We confirmed the existence of some common chimeric RNAs and also discovered many novel chimeric RNAs during retinal development. We focused on CTNNBIP1-CLSTN1 (CTCL) whose downregulation causes precocious neuronal differentiation and a marked reduction of neural progenitors in human cerebral organoids. Our study found that CTCL also plays a key role in human retinogenesis, CTCL loss-of-function obstructed RO differentiation but prompted the retinal pigment epithelial (RPE) differentiation. Together, this work provides a landscape of chimeric RNAs and reveals evidence for their crucial roles in human retina development.

## Introduction

The human retina is a laminal structure with a large number of different constituent cells that form morphologically and functionally distinct circuits. They work in parallel and in combination to produce complex visual output (Hoon et al., 2014). During retinogenesis, different subtypes of neurons are generated from the same group of retinal progenitor cells and self-assembled accurately into a functionally mature retina (Masland, 2012).

Dissecting the molecular mechanisms of human retinogenesis and functional maintenance has always been the main focus and a difficult issue, which is particularly important for the treatment of human-specific diseases such as macular degeneration. A few groups systematically expounded the transcriptomics, chromatin accessibility and proteomic dynamics during human and mouse retinogenesis (Huang et al., 2022), and comprehensively described the similarities and differences during this process. They identified an unexpected role for ATOH7 expression in regulation of photoreceptor specification during late retinogenesis (Lu et al., 2020), as well as the enriched bivalent modification of H3K4me3 and H3K27me3 in human but not in murine retinogenesis (Lu et al., 2020), further underscore the limitations of using the mouse models to study human retinal and the critical role of transcriptional regulation in human retinogenesis.

Chimeric RNAs are the ligation products of two or even more DNA or RNA sequences before or after transcription. They can be further translated into proteins in addition to their RNA form, which increases the richness of the transcriptome and proteome. Previous studies of chimeric RNAs have focused on tumors and larger tissues and organs, and found that chimeric RNAs are widely present in human normal and tumor tissues as part of the transcriptome and can regulate the life activities of individual cells (Elfman & Li, 2018; Hu et al., 2018; Mertens et al., 2015; Singh et al., 2020). It has been shown that chimeric RNAs in normal human cortex increase with age and show differences in individual cells and tissues (Mehani et al., 2020), which highlights the critical role of chimeric RNAs in CNS development and cell lineage maintenance. In 2021, Luo’s group found that downregulation of CTCL affects cerebral organoids growth, causing premature neuronal differentiation and a marked reduction of neural progenitors (Ou et al., 2021). As retina is a part of the CNS, we hypothesize that CTCL also exists in the retina and plays an important role in retinal development.

The role of chimeric RNAs in human retinogenesis remains unexplored. This study aims to investigate the molecular mechanism of human retinal development and function maintenance in a new perspective of chimeric RNAs. Difficulties in obtaining normal human retinal tissue have hampered studies related to human retinal development. The emergence of ROs could circumvent this problem (Jin et al., 2019). Our group has extensive experience in retinal organoid culture and has conducted a series of studies using ROs (Deng et al., 2018; Ma et al., 2022; Pan et al., 2020; Zhang et al., 2021). For example, we, along with other groups, used ROs to confirm that retinoblastoma tumor cells are derived from retinal maturing cone cells (Clevers, 2016; Jin et al., 2019; Li et al., 2022; Liu et al., 2021; Liu et al., 2020).

In this study, we use hESC-derived ROs and perform RNA sequencing. We found that chimeric RNA CTCL is critical in the process of organoid differentiation towards ROs or RPE. Retinal progenitor cells with a relatively high level of CTCL expression will differentiate towards ROs, while those with a relatively low level of CTCL will differentiate towards RPE. These results provide important evidence for the physiological function of chimeric RNAs in retinogenesis.

## Results

### Chimeric RNAs are present throughout the retinogenesis

To explore the chimeric RNAs in the developing human retina, we cultured ROs according to the methods described previously and collected organoid tissues from 0 to 120 days for RNA sequencing (Kim et al., 2019; Lowe et al., 2016) (Figure 1, top panel and middle panel). Using FusionCatcher software, we examined chimeric RNAs in 22 human ROs covering eight developmental stages, including day 0 (D0), 30, 45, 60, 75, 90, 105 and 120, corresponding to in vivo developmental stages D0, 30, 45, 60, 75, 90, 105 and 120 (Cowan et al., 2020). To reduce false positive results, only if spanning unique reads were recognized at least once, the splicing events would be considered as positive chimeric RNA candidates (Figure 1, bottom panel).

**Figure 1.**
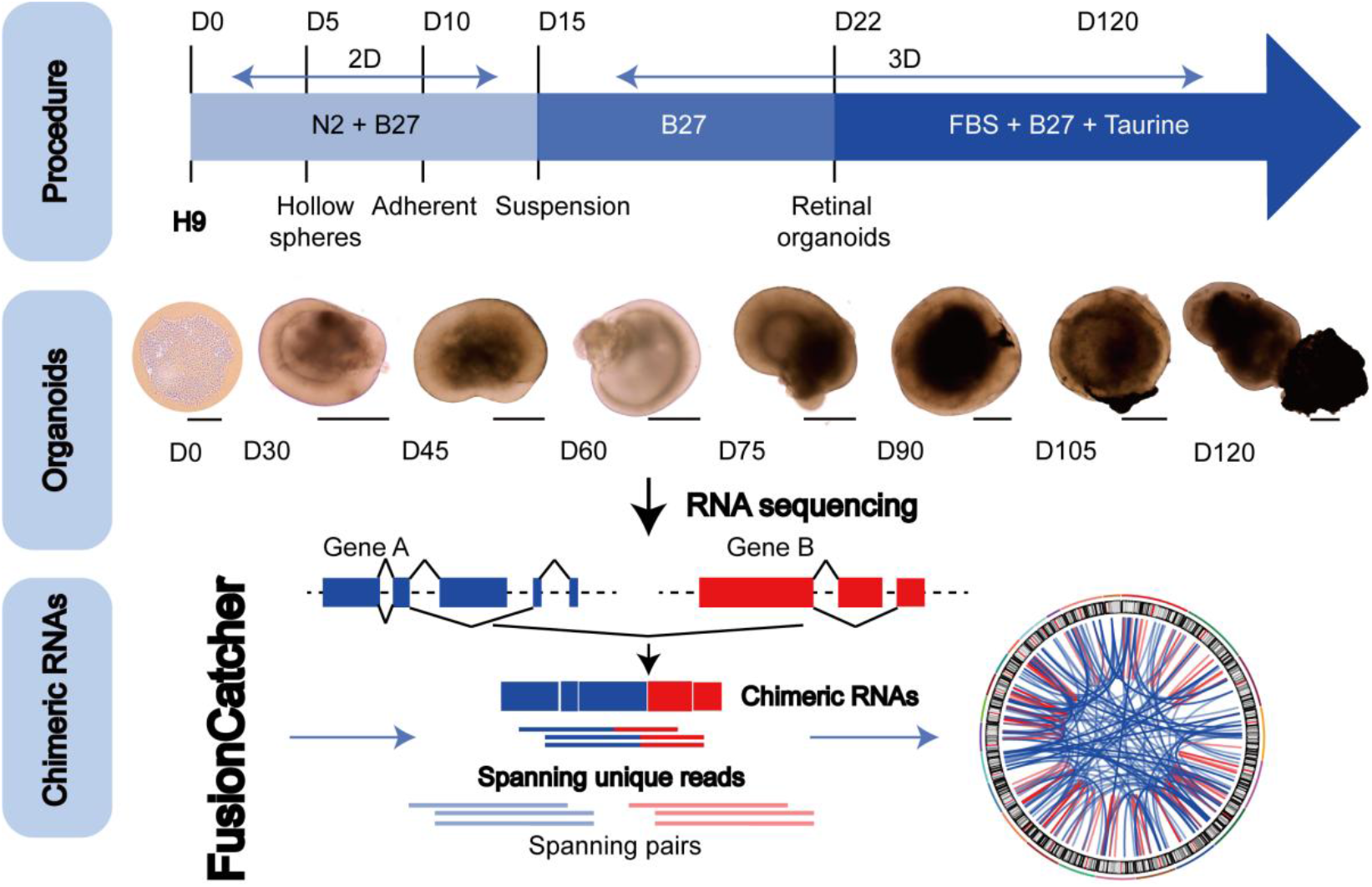
The general scheme of detection of chimeric RNAs. (Top panel) Procedure for the generation of human ROs. (Middle panel) Human ROs of 0-120 days were used for sequencing, scale bars: 200 μ m (D0) and 400 μ m (D30-D120). (Bottom panel) Illustration of chimeric RNAs screening process and criteria by FusionCatcher.

Chimeric RNAs were continuously expressed during RO development (Figure 2A). Chimeric events were categorized according to the locations of the parental genes on the chromosome, either inter-chromosomal or intra-chromosomal. While the number of intra-chromosomal chimeric RNAs increased with the development of ROs (*r*^*2*^ = 0.93, *p* = 0.00076, Pearson’s correlation analysis), there was no such trend for inter-chromosomal chimeric RNAs (Figure 2B-C). Chimeric events can also be categorized according to the predicted effect (Figure 3–figure supplement 1). In the top 3 categories, the number of “In frame” chimeric RNAs was 11.2%, followed by “CDS(truncated)_UTR” (10.7%) and “UTR_UTR” (10.7%) (Figure 3A). We further classified chimeric events according to the type of their parental genes. The majority of chimeric RNAs were formed between two protein-coding genes (61.9%) (Figure 3B).

**Figure 2.**
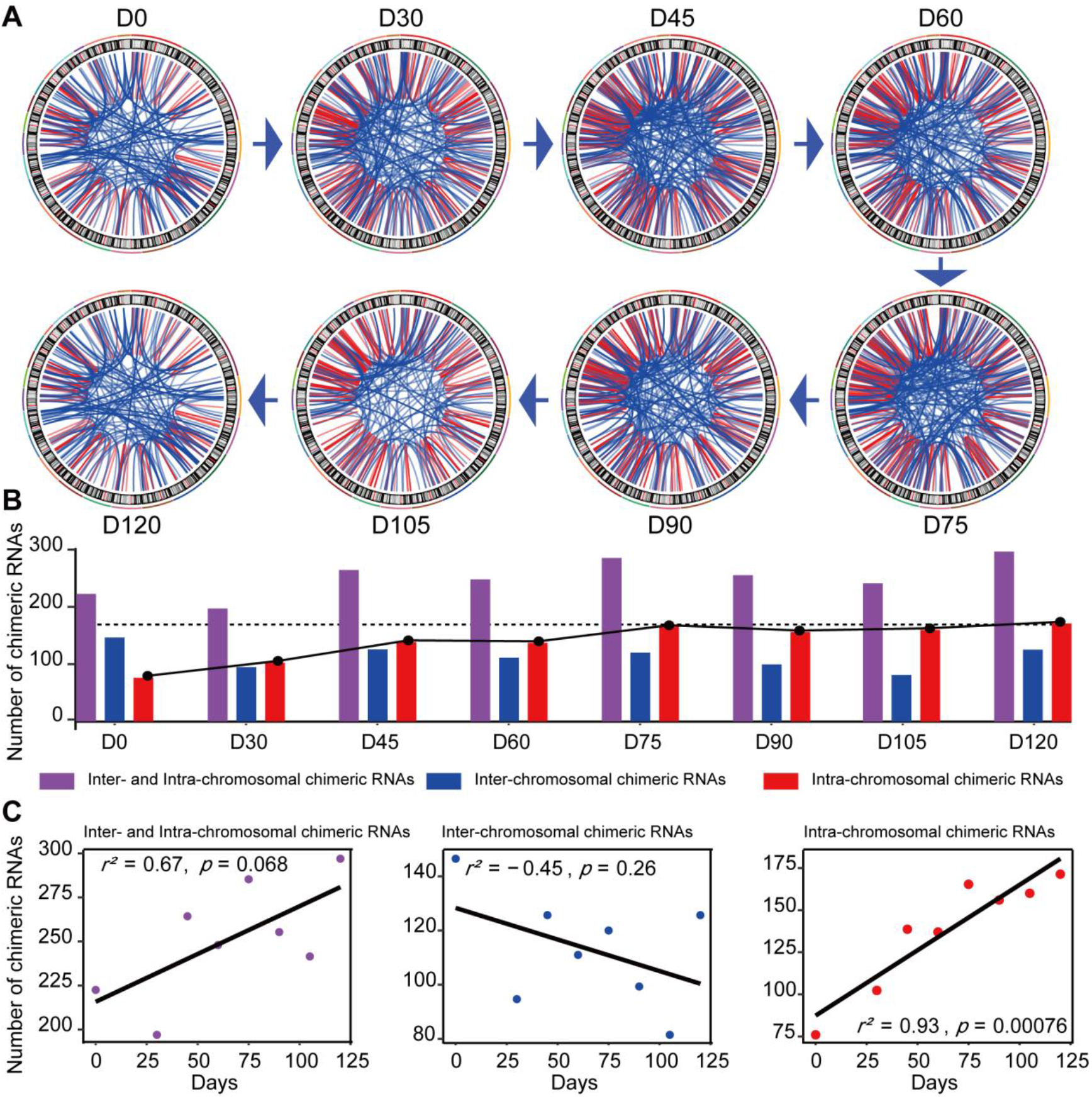
Expression of chimeric RNAs in the developing human retinal organoids. **(A)** Circos plots of genomic distribution of chimeric RNA parental genes observed in this work. Red lines indicate parental genes located in the same chromosome. Blue lines indicate parental genes located in different chromosomes. The outermost colored lines of circos plots represent chromosomes. **(B)** Types of chimeric RNAs based on parental genes’ genomic distribution. **(C)** Correlation analysis of the number of chimeric RNAs and developmental stages with Pearson’s correlation analysis.

**Figure 3.**
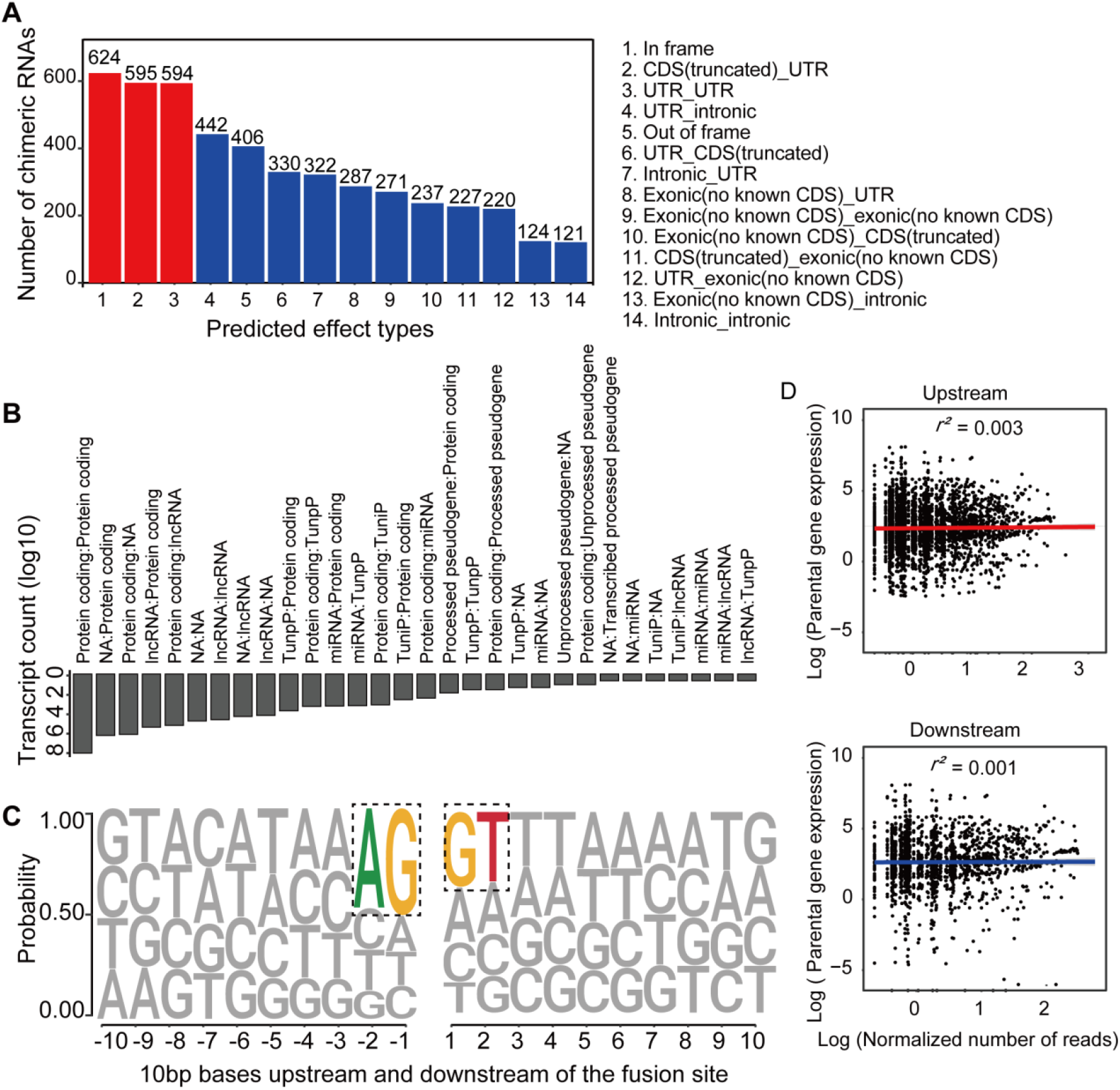
Characterization of chimeric RNAs in the developing human retinal organoids. **(A)** Top 14 types and corresponding numbers of chimeric RNAs based on predicted effects. **(B)** Biotype quantification of parental gene combinations in all samples. TunpP: Transcribed unprocessed pseudogene; TuniP: Transcribed unitary pseudogene. **(C)** Motifs consisting of 20 bp DNA sequences around the fusion site. **(D)** Spearman correlation analysis of expression level of chimeric RNAs and their parental genes, *p* > 0.05.

Next, we examined the motifs covering 10-bp sequences immediately upstream or downstream to the fusion site of parental genes and found that the canonical GT/AG donor-acceptor motif had the highest position weight (Figure 3C). In addition, we also found that the expression of chimeric RNAs did not correlate with the expression of parental genes (Figure 3D). These results were consistent with previous reports (Ou et al., 2021; Singh et al., 2020), indicating that the procedure of screening chimeric RNAs was reliable and those chimeric RNAs do not arise randomly, which suggested that they play a role in the normal development and physiological activity of the retina.

### CTCL is present in all the stages of human retinal development

CTCL has been shown to have a regulatory role for human cerebral development. Similarly, four isoforms of CTCL joined by alternative splicing of parent pre-mRNAs are detected in the ROs RNA-seq results. In-frame CTCL was dynamically expressed during RO development (Figure 4A-B) and there are two obvious higher expression timepoints, D60 and D120, respectively. Around D120, retinal cells like cones, rods, horizonal cells have been generated, therefore, we chose D60 when most retinal cells are progenitor cells. To further validate the presence of CTCL in the developing human ROs, we extracted total RNAs from D60 human ROs, followed by reverse transcription and Sanger sequencing using specific primers to amplify the fragments containing fusion sites between CTNNBIP1 and CLSTN1 (Figures 4C-D and table S1). At D60, in-frame, UTR_CDS(truncated)-1 and CDS(truncated)Intronic CTCL isoforms were confirmed but UTR_CDS(truncated)-2 CTCL was not detected. The dynamic presence of CTCL, especially the in-frame isoform, suggested that it may play a role in human retina development.

**Figure 4.**
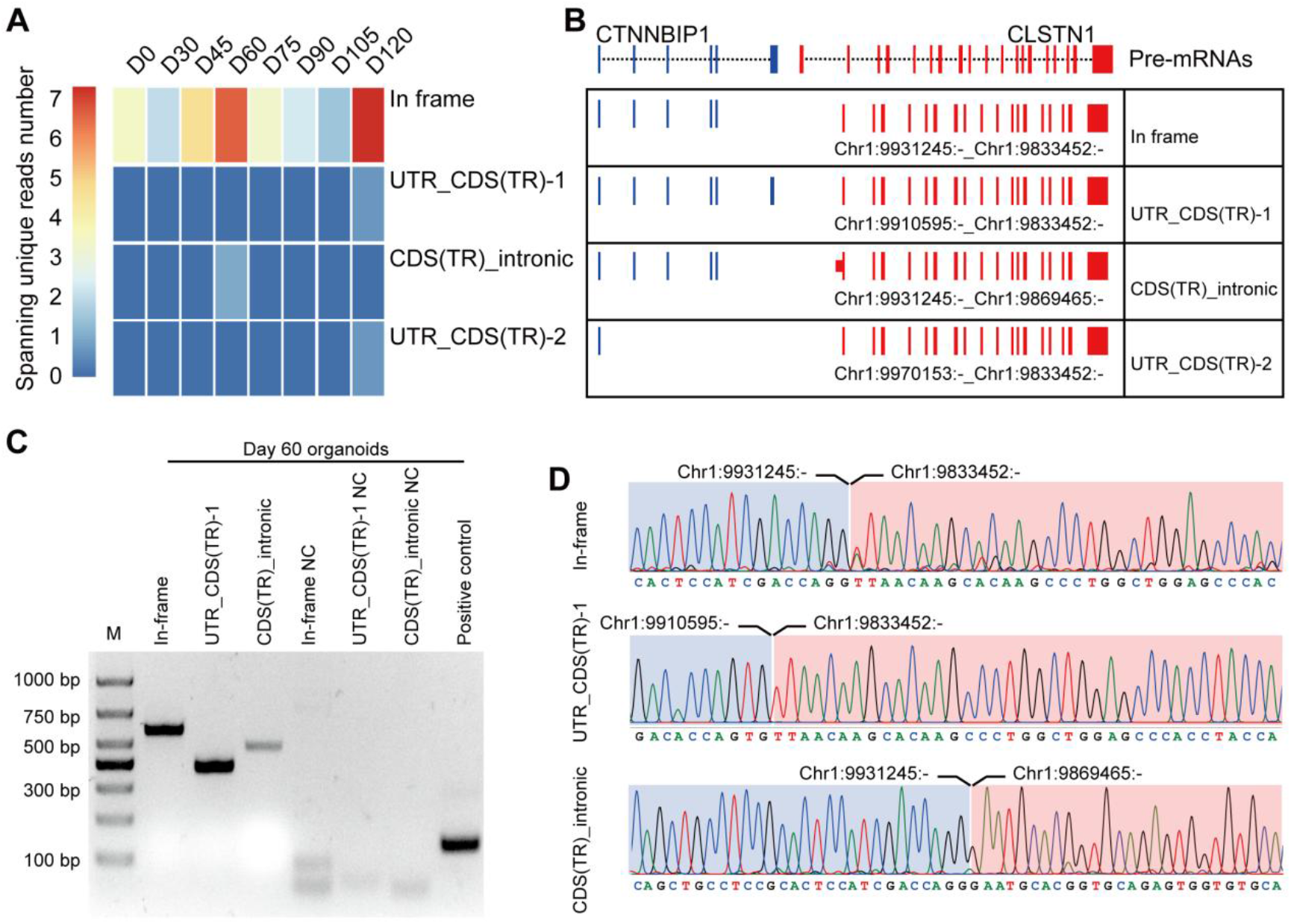
Four isoforms of CTCL are present in the retinal organoids. **(A)** Heatmap of CTCL’s spanning unique reads of each isoform in the indicated stages. UTR_CDS(TR)-1: UTR_CDS(truncated)-1; CDS(TR)Intronic: CDS(truncated)Intronic; UTR_CDS(TR)-2: UTR_CDS(truncated)-2. **(B)** Schematic diagram of the structures of the four isoforms of CTCL. Blue represents upstream parental gene, red represents downstream parental gene. **(C)** PCR validation of four isoforms of CTCL in ROs at day 60. **(D)** Sanger sequencing to verify three isoforms of CTCL.

### The CTCL knockdown obstructed ROs differentiation but prompted the RPE differentiation

According to the CTCL RNA expression levels (Figure 4A), we focus on the D60 in-frame CTCL to explore the function of chimeric CTCL. A hPSC line with CRX-tdTomato reporter was used in the CTCL knockdown experiment (Pan et al., 2020). ROs were differentiated according to the previous report (Lowe et al., 2016) (Figure 5 – figure supplement 1) (Video 1). Lentivirus carrying shCTCL (short-hairpin CTCL) or scramble shRNA (Table S1) infected the retinal cells at D12. After 3 days, cells were collected for the knockdown efficiency analysis (Figure 5A-B). Comparing with the scramble shRNA group, in-frame CTCL RNA level was reduced to around 70% (Figure 5B). The RNA levels of parental genes, *CTNNBIP1* and *CLSTN1*, displayed no significant differences in comparison with the scramble shRNA group (Figure 5–figure supplement 2).

**Figure 5.**
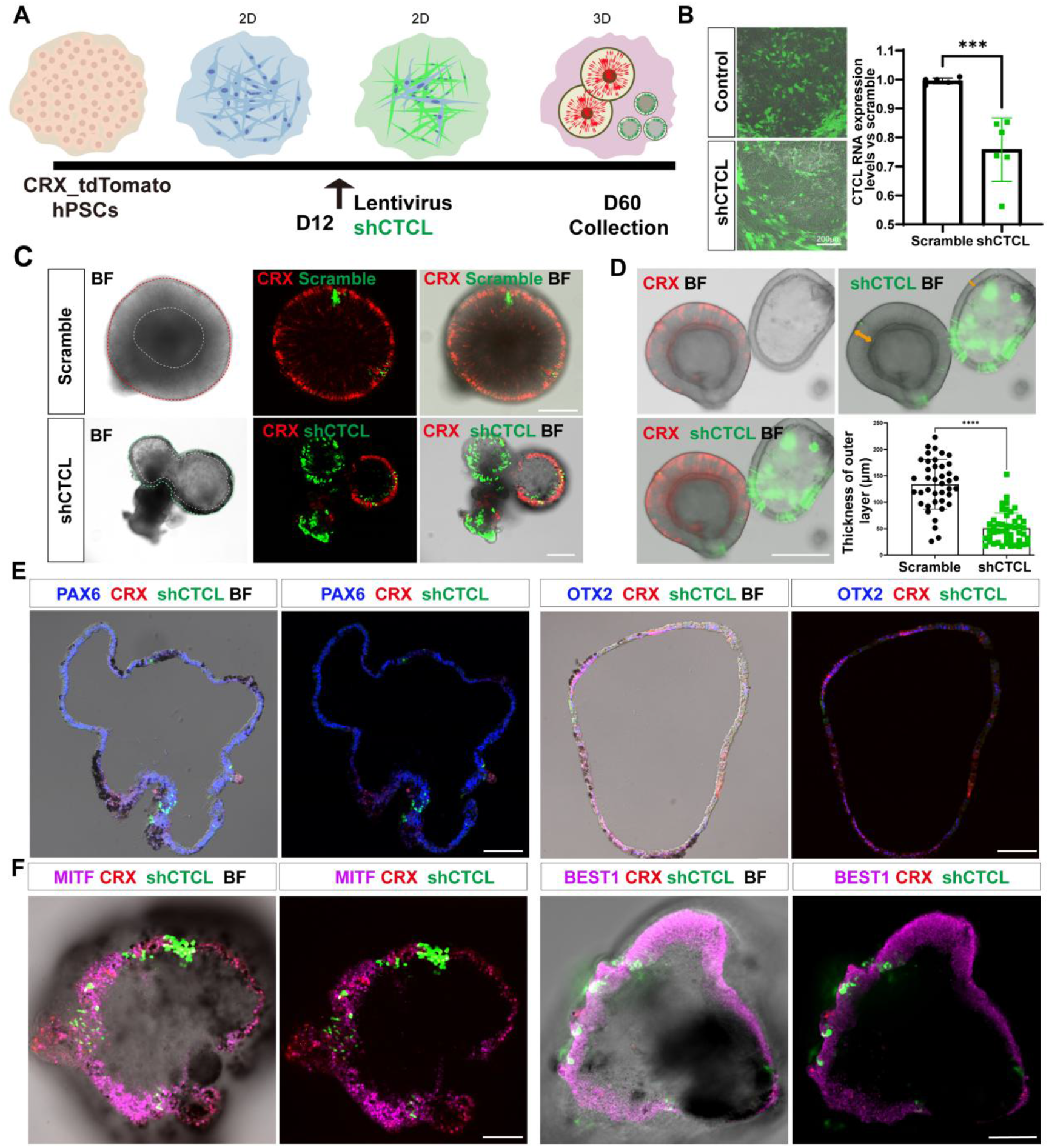
The CTCL knockdown obstructed ROs’ differentiation but prompted the RPE differentiation. **(A)** A schema illustrated the shRNA experiments. A CRX-tdTomato report line was used in this experiment. The shCTCL or scramble shRNA lentivirus transfected the retinal cells in three independent experiments on D12. All samples for shRNA experiments were collected on day 60, except those for analysis of knockdown efficiency. **(B)** Samples were obtained 48-72 hours after infection to examine the knockdown efficiency. There were two technical replicates for three independent experiments, *t-test, p* < 0.001. **(C)** D60 shRNA (Scramble)-transfected ROs displayed a typical morphology of ROs with the expression of CRX. ShCTCL-transfected organoids showed thinner outer layers with much less CRX expression. BF, bright-field images, same as below. **(D)** The shCTCL group displayed much thinner outer layers (orange arrows in right panel). The outer layer thickness of 25 organoids from each group was measured for statistics analysis, *t-test, p* < 0.0001. **(E)** The section immunostaining of neural progenitor markers PAX6 and OTX2, and differentiated cell markers CRX, in shCTCL-treated organoids. Scale bars = 100μm. **(F)** Whole-mount immunostaining of shCTCL-transfected organoids with CRX, shCTCL, and RPE specific markers MITF and BEST1. Scale bars = 100μm.

At D60, ROs with scramble shRNA displayed the typical RO morphology with the generation of photoreceptor precursors (CRX^+^) (Figure 5C, upper panel). In the shCTCL group, the morphology of the organoids changed with a thinner outer layer, and the CRX proteins diminished where the pigments appeared (Figure 5C, lower panel). We measured the average thickness of the outer layers in those two groups (Figure 5D, outer layer indicated with orange arrows), there is a dramatic decrease of the outer layer thickness in the shCTCL group (avg±sd vs avg ±sd).

In order to identify the aberrations in cell fate specification caused by CTCL knockdown, we examined the organoids with retinal progenitor specific markers, PAX6 and OTX2. It showed that PAX6 was widely expressed, but OTX2 was much less, and CRX appeared when shCTCL was low (Figure 5E). As we know, CRX is highly expressed in photoreceptors and lowly expressed in mature RPE (Lidgerwood et al., 2021). While the organoids acquired the features of RPE (pigments), we believed that the outer layer of CTCL-knockdown organoids switched to RPE fates. To further confirm this, we were immunostained the organoids with RPE-specific markers, MITF, BEST1 and RPE65, which were present in the nucleus, cytoplasm and intracellular regions, respectively (Figure 5F, Figure 5–figure supplement 3A, Figure 5–figure supplement 3C). By contrast, no pigments or RPE-specific markers were observed in the scramble shRNA group, only the PAX6 was detected in the nucleus (Figure 5–figure supplement 3B, Figure 5–figure supplement 3D-F). These results indicated that CTCL is essential for the ROs’ differentiation and CTCL knockdown would promote the RPE cell fate at the expense of neural retinal cells.

### The underlying molecular changes in CTCL knockdown ROs

The above results suggested that CTCL is essential for cell fate conversion in normal retinogenesis. To investigate the underlying molecular regulation, we compared the transcriptomes of shCTCL and scramble shRNA-treated organoids by analyzing the bulk RNA-seq results. We obtained 3901 differentially expressed genes (DEGs) (shCTCL vs. scramble shRNA, |Log2 (fold change)| > 1, p-value < 0.05), among which 2720 were upregulated and 1181 down regulated (Figure 6A-B).

**Figure 6.**
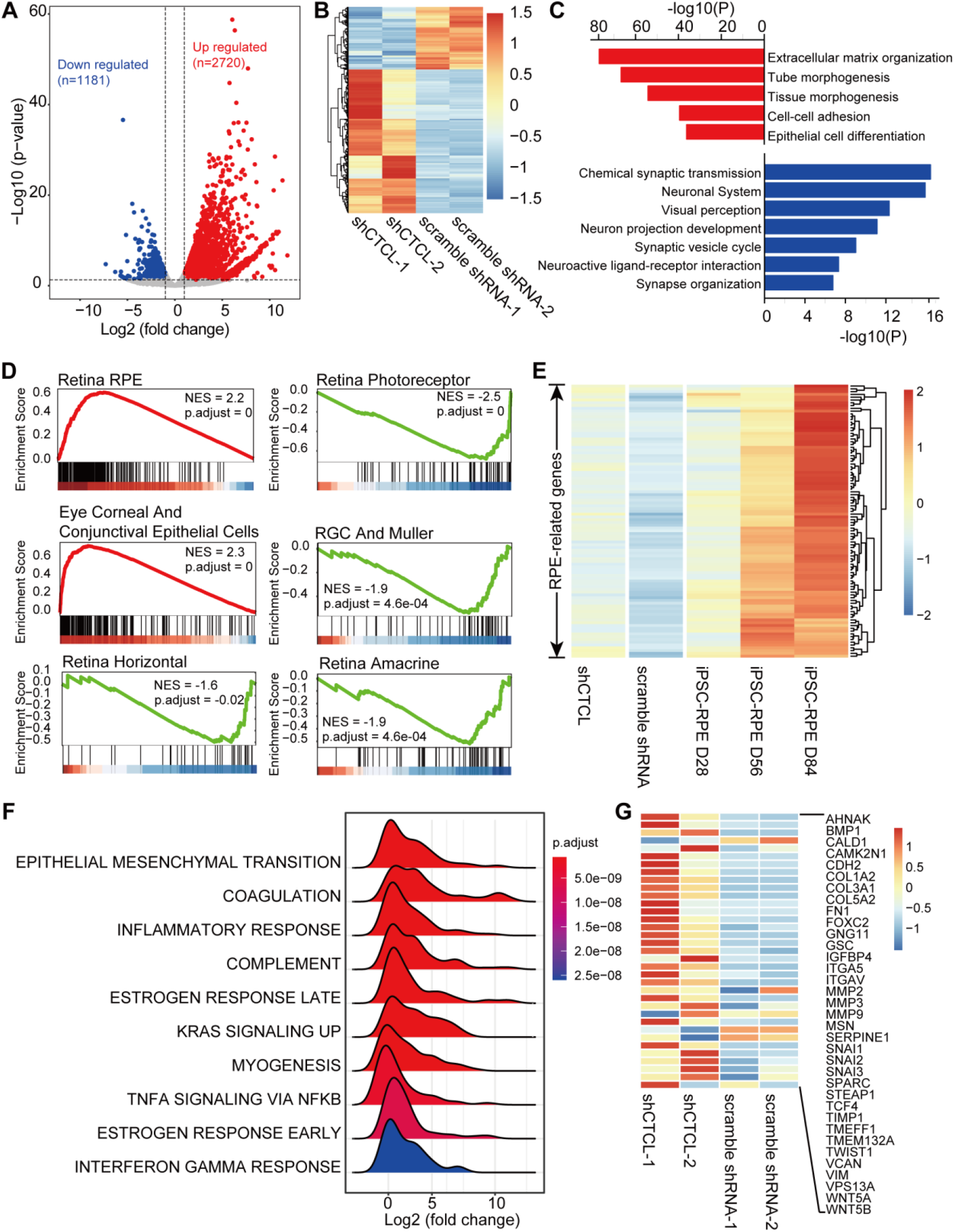
Transcriptome alterations in CTCL-knockdown retinal organoids. **(A)** Volcano plot shows differentially expressed genes (p-values < 0.05 and |Log2 (shCTCL/scramble)| > 1) between shCTCL and scramble shRNA-treated organoids at D60. **(B)** Heatmap of DEGs in shCTCL and scramble shRNA-treated organoids. **(C)** Functional enrichment analysis of up- and down-regulated DEGs. **(D)** GSEA results showed enriched gene sets in shCTCL-treated organoids using the c8 reference gene set. **(E)** Expression of RPE-related genes in shCTCL-treated, scramble shRNA-treated organoids. **(F)** GSEA results showed the enriched gene sets in shCTCL-treated organoids using the hallmark reference gene set. **(G)** Expression of EMT related genes in shCTCL-treated, scramble shRNA-treated organoids.

To investigate the biological pathways affected by the loss of function of CTCL, we used Metascape to perform functional enrichment for upregulated and downregulated DEGs. Upregulated DEGs are significantly enriched in the extracellular matrix organization, morphogenesis and epithelial cell differentiation associated pathways. On the other hand, downregulated DEGs are significantly enriched in the neurogenesis associated pathways (Figure 6C). To further understand the changes in cell composition between shCTCL and scramble shRNA-treated organoids, we used the Gene Set Enrichment Analysis (GSEA) to analyze whole gene list of shRNA-treated organoids. GSEA analysis with c8 as a reference gene set (Mootha et al., 2003; Subramanian et al., 2005) revealed that genes of RPE, eye corneal and conjunctival epithelial cells were significantly upregulated with shCTCL organoids; however, all genes related to neural retinal component cells (photoreceptor cells, mü ller glial cells, horizontal cell, amacrine cell, retinal ganglion cell) were downregulated (Figure 6D). Moreover, the expression pattern of RPE-related genes in shCTCL organoids was similar to that of RPE cultured from induced pluripotent stem cells (Maruotti et al., 2015) (Figure 6E and Figure 6–figure supplement 1A). At the same time, CTCL maintains at a low expression status during RPE development (Figure 6–figure supplement 1B). These results suggested that CTCL has a positive role in the normal development of ROs and that CTCL deficiency would promote the differentiation of retinal progenitor towards RPE.

To further explore the molecular mechanisms involved in this process, we used hallmark gene sets (Mootha et al., 2003; Subramanian et al., 2005) to perform GSEA analysis. The most significantly enriched pathway was epithelial mesenchymal transition (EMT) (Figure 6F). And most EMT-related genes are relatively highly expressed in shCTCL-treated organoids (Figure 6G). which is particularly essential for neural crest delamination of vertebrates and for generating different tissues during organismal development (Kim et al., 2014). Therefore, we believe that the downregulation of CTCL expression activated the EMT pathway which altered the directions of cell differentiation.

## Discussion

The term “chimeric RNA” refers to any transcript consisting of transcripts from different parental genes, including gene fusion transcripts (Wu et al., 2019). Chimeric RNAs are formed by splicing transcripts from two parental genes, a process that generates new dysregulated wild-type proteins, new fusion proteins and new non-coding RNAs, expanding the abundance of transcriptome and proteome. The first chimeric RNA was discovered to be caused by chromosomal rearrangements, i.e., BCR-ABL1, which is associated with chronic leukemia (Ren, 2005). Subsequently, cis-splicing between neighboring genes was also found to generate chimeric RNAs, such as CTCF-sensitive cis-spliced fusion RNAs that can cause prostate cancer (Qin et al., 2015). Addition, chimeric RNAs can be generated by long-distance inter- and intra-chromosomal trans-splicing (Li et al., 2008). Early phase of studies had focused on the role of chimeric RNAs in cancer, where they contribute to cancer development, and serve as biomarkers and therapeutic targets.

It is now well established that chimeric RNAs occur abundantly in normal human tissues and have a regulatory role in cellular life activities. In this work, we study chimeric RNAs in retina for the first time. We present the expression atlas of chimeric RNAs throughout the developing ROs. We identified three isoforms of CTCL, and the in-frame isoform has been reported to play a key role in cerebral development previously (Ou et al., 2021). By using the ROs as model for loss-of-function experiments, we found that the in-frame CTCL also has a key role in the development of the retina and that CTCL deficiency obstructed RO differentiation but prompted the RPE differentiation.

In-frame CTCL is generated by cis-splicing fusion of the first 5 exons of CTNNBIP1 and the last 17 exons of CLSTN1. The 5’ parental gene CTNNBIP1 is a receptor for β-catenin, and their binding promotes the catabolism of β-catenin, which in turn puts the wnt pathway in an inactive state (Fu et al., 2018). CLSTN1 is a member of the calsyntenin family, a subset of the cadherin superfamily. It can mediate the axonal anterograde transport of certain types of vesicles (Nagase et al., 1998). The interesting question is that whether CTCL keeps the biparental functions and/or acquired novel functions? Previous report demonstrated that CTCL could fine-tune wnt signaling to regulate cerebral development (Ou et al., 2021). Therefore, in this study, we had expected that CTCL functions by modulating the wnt signaling pathway or by impacting vesicular transport. However, surprisingly, the GSEA and DEG functional enrichment results did not show significant changes in these two biological pathways (Figure6 C, D, F). The differences between the shCTCL or scramble shRNA-treated organoids were mainly about EMT, extracellular matrix organization, morphogenesis, epithelial cell differentiation and neurogenesis associated pathways, which implicate that even the same chimeric RNAs may function differently in different tissues.

Chimeric RNAs can function in both transcriptome and proteome levels. In this study, we verified CTCL expression at the transcriptional level but failed to detect its protein as in brain organoids (Ou et al., 2021). The differentiation of RPE from human parthenogenetic embryonic stem cell was previously reported and miR-204 targeting CTNNBIP1 was found to determine this process (Li et al., 2012). This suggests that regulation at the RNA level can play a decisive role in the differentiation of RPE. In our study, we found that the expression levels of both parent genes (CTNNBIP1 and CLSTN1) and CTCL were not correlated, and that the expression levels of both parent genes were unaffected between shCTCL or scramble shRNA-treated organoids. Therefore, we suggest that CTCL exerts its regulatory roles in human retinal development mainly through RNA rather than protein, which may also explain why CTCL affects different biological pathways in cerebral and retinal development.

In summary, we identified the chimeric RNAs in the developing human ROs and found that loss-of-function of CTCL obstructed neuroretinal differentiation while prompted the RPE differentiation. This study for the first time revealed a key role of chimeric RNAs in human retinal development, providing new insights of chimeric RNAs in regulating tissue development.

## Materials and methods

### RNA sequencing and data analysis

A total amount of 1-3μg RNA per sample was used as input material for the RNA sample preparations. Sequencing libraries were generated using VAHTS Universal V6 RNA-seq Library Prep Kit for Illumina ® (NR604-01/02) following the manufacturer’s recommendations and index codes were added to attribute sequences to each sample. Briefly, mRNA was purified from total RNA using poly-T oligo-attached magnetic beads. Then we added fragmentation buffer to break the mRNA into short fragments. First strand cDNA was synthesized using random hexamer primer and RNase H. Second strand cDNA synthesis was subsequently performed using buffer, dNTPs, DNA polymerase I and RNase H. And then, the double stranded cDNA was purified by AMPure P beads or QiaQuick PCR kit. The purified double stranded cDNA was repaired at the end, added a tail and connected to the sequencing connector, then the fragment size was selected, and finally the final cDNA library was obtained by PCR enrichment.

We used FusionCatcher software (https://github.com/ndaniel/fusioncatcher) to identify chimeric RNAs in human ROs. Positive chimeric RNAs identified using FusionCatcher were selected with alignment of spanning unique reads. The expression level of chimeric RNAs was reflected from log (Spanning_unique_reads/Total_number_of_reads*10000000) value. To analyze the expression of parental genes in human ROs, raw reads were first mapped to the hg38 human genome reference sequence by Hisat2 software, then transcripts were assembled with featureCounts. Metascape (https://metascape.org/gp/index.html) is used for functional annotation. The position weight matrix of the 20 bp DNA sequence motif around the fusion site was calculated by the seqLogo R package. GSEA analysis is done with the R package clusterProfiler.

### Generation of ROs from hESCs

The CRX-tdTomato human ES reporter line was used for the ROs differentiation. The hESCs were cultured to 80% confluence. Cell colonies were dissociated using dispase buffer (Stem cell, 07923) for 5 minutes at 37 °C and were then cut into smaller pieces/aggregates. The aggregates were collected and mixed with Matrigel (RD). The solidified gel with ES cells were dispersed into small pieces with medium containing 1:1 mixture of DMEM/F12, neurobasal medium, and 0.5x N2 supplement (GIBCO), 0.5x B27 supplement (GIBCO), 1x MEM-NEAA (GIBCO), 2 mM Glutamax (GIBCO), 0.1 mM 2-mercaptoenthanol. At day 1 (D1), hollow cysts could be observed in the dish and cysts started to attach to the culture dish and spread in 3 days. At D5, the cysts were dispersed into two culture dishes with same medium. Medium was changed every 5 days. On D15, detaching the cells with dispase and change to medium with DMEM/F12 (3:1), 1x B27 supplement (GIBCO), 1x MEM-NEAA (GIBCO) for a week. Optic vesicles formed in this period. Finally, optic vesicles were transfered to serum medium with DMEM/F12 (3:1), 1x B27 supplement (GIBCO), 1x MEM-NEAA (GIBCO), 8% FBS (GIBCO), 100 mM Taurine (Sigma-Aldrich) and 2 mM Glutamax (GIBCO). ROs would be collected at D60.

### CTCL knockdown in the ROs

RNA interference oligo (5’-TGCTTGTTAACCTGGTCGA-3’) against CTCL was cloned into lentivirus vector (pLenti-U6-shRNA-EF1a-EGFP-T2A-Puro-WPRE). D12 retinal cells were infected with lentiviruses (MOI=10) for 6 hours, and replaced with fresh medium. The fluorescence was detected after 48 hours. The expression of CTCL, CTNNBIP1 and CLSTN1 in retinal cells were analyzed at D15 by quantitative PCR.

### Immunostaining of cryosections

ROs were fixed in 4% PFA for 30 minutes and imbedded in O.C.T. compound and sectioned into 10 μm slices. The cryosections were blocked with 0.5% Triton X-100 in 4% BSA for 1 hour. After that, sections were incubated in primary antibodies (diluted in 4% BSA supplied with 0.5% Triton X-100) at 4°C overnight. The following primary antibodies were used: anti-OTX2 (1:200, Cat. #ab183951; Abcam), anti-PAX6 (1:200, Cat.# 901301;Biolegend), anti-SOX2(1:200, Cat.# sc-365823; Santa Cruz), anti-HuC/D (1:100, Cat.# A21271; Invitrogen), anti-MITF (1:100, Cat. # ab3201; Abcam), anti-GFAP (1:200, Cat.# sc-33673; Santa Cruz), anti-Sox9 (1:200, Cat.# 711048; Invitrogen), anti-RxRγ (1:100, Cat.# sc-365252; Santa Cruz), and anti-Ki67 (1:200, Cat.# ab15580; Abcam). After washed with PBS, cryosections were stained with Alexa Fluor-conjugated secondary antibodies (diluted 1:500, Invitrogen) for 1 hour at room temperature in the dark.

### Quantification and statistical analysis

Heatmap was plotted with the R package Pheatmap. All plots were drawn with R package ggplot2. Detailed statistical analysis of experiments can be found in the figure legends, including the statistical tests used, exact values and biological replicates.

## RESOURCE AVAILABILITY

Further information and requests for reagents should be directed to and will be fulfilled by the lead contact, Professor Zi-Bing Jin.

## Materials availability

All unique/stable reagents generated in this study are available from the lead contact with a completed materials transfer agreement.

## Data and code availability

The Gene Expression Omnibus (GEO) numbers for the bulk RNA-seq of organoids in this paper are GEO: GSE136929. All software used is open and freely available. The published article includes main datasets generated during this study.

## Acknowledgments

This study was partially supported by grants from National Natural Science Foundation of China (82125007).

## Author contributions

Zi-Bing Jin designed and supervised the study, provided financial supports and revised the manuscript; Wen Wang performed the computational analysis and interpreted the data, wrote the manuscript; Xiao Zhang carried out the experiments; Ning Zhao and Ze-Hua Xu performed the stem cell differentiation; Kangxin Jin interpreted the results and revised the manuscript.

## Video and supplementary

**Video 1**. The 4 days live cell imaging of retinal organoid differentiation from D3 to D7.

**Supplementary table 1**.Key resource tables.

**Figure 3–figure supplement 1.**
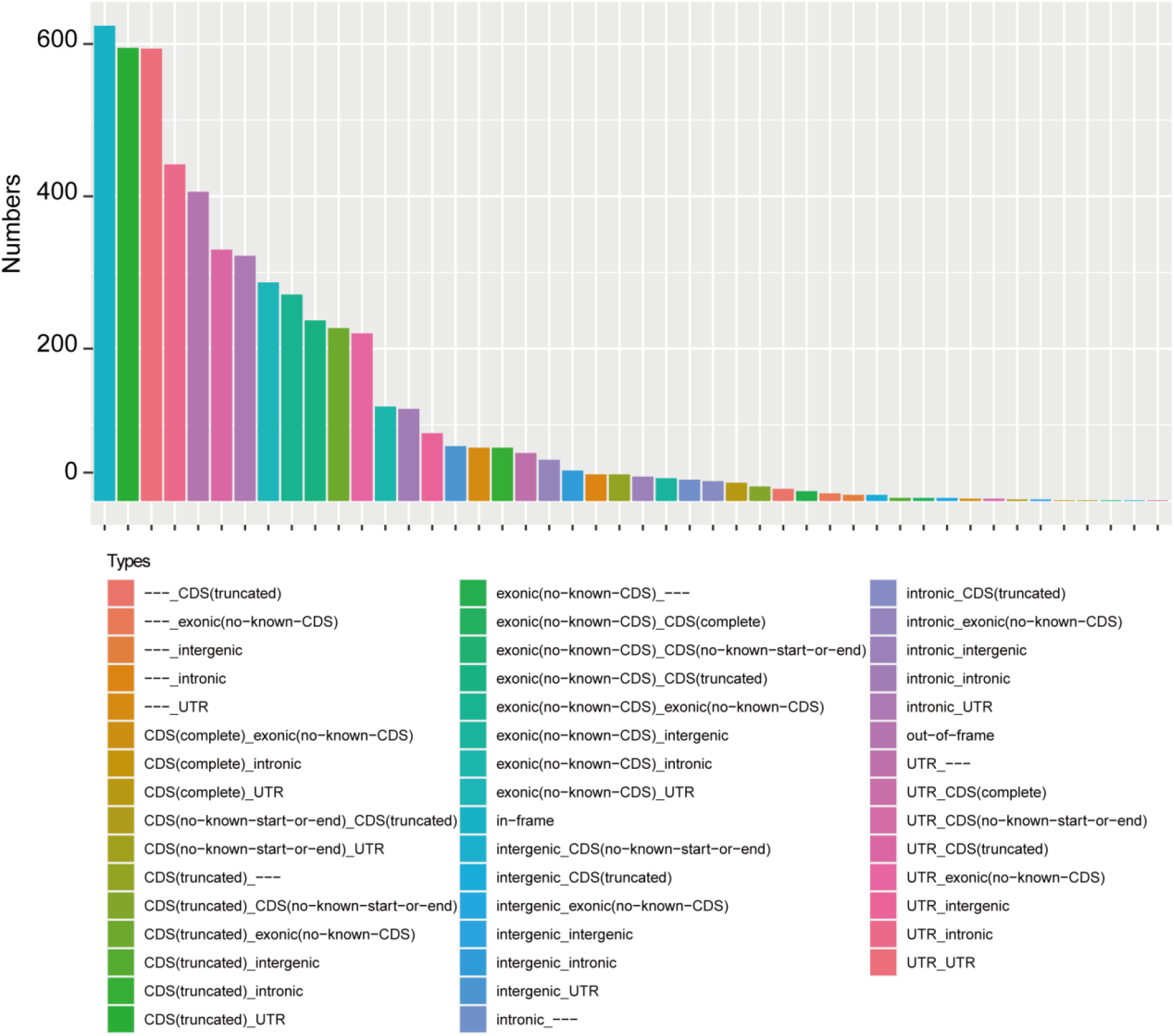
Types of chimeric RNAs based on predicted effect.

**Figure 5–figure supplement 1.**
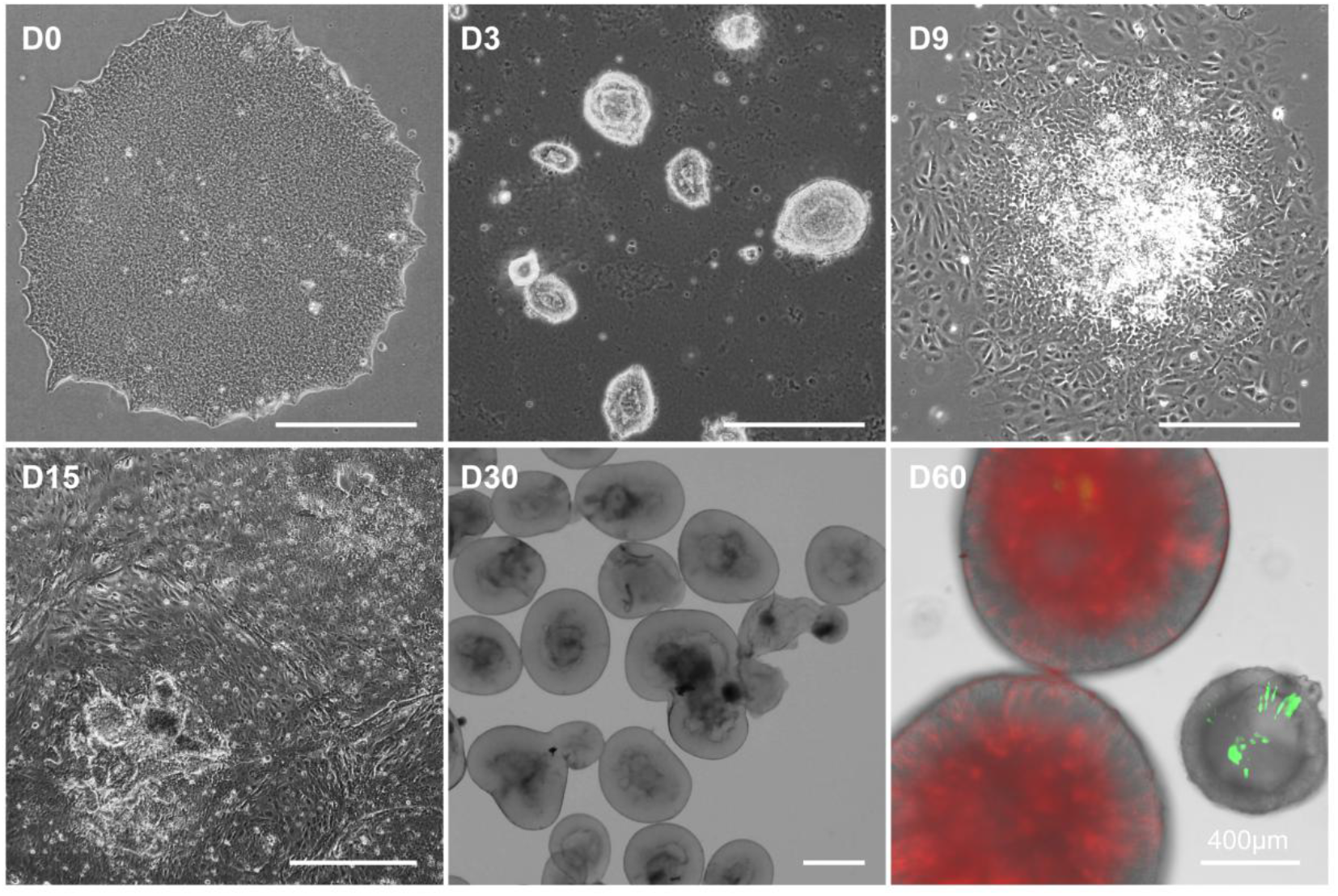
Images at different timepoints. This RO differentiation method was a combination of 2D and 3D culture. On D60, CTCL downregulated organoids displayed a thinner outer layer. Live imaging showed D60 organoids were represent with CRX reporter tdTomato (red) and shCTCL reporter GFP (green). Scale bars = 400μm.

**Figure 5–figure supplement 2.**
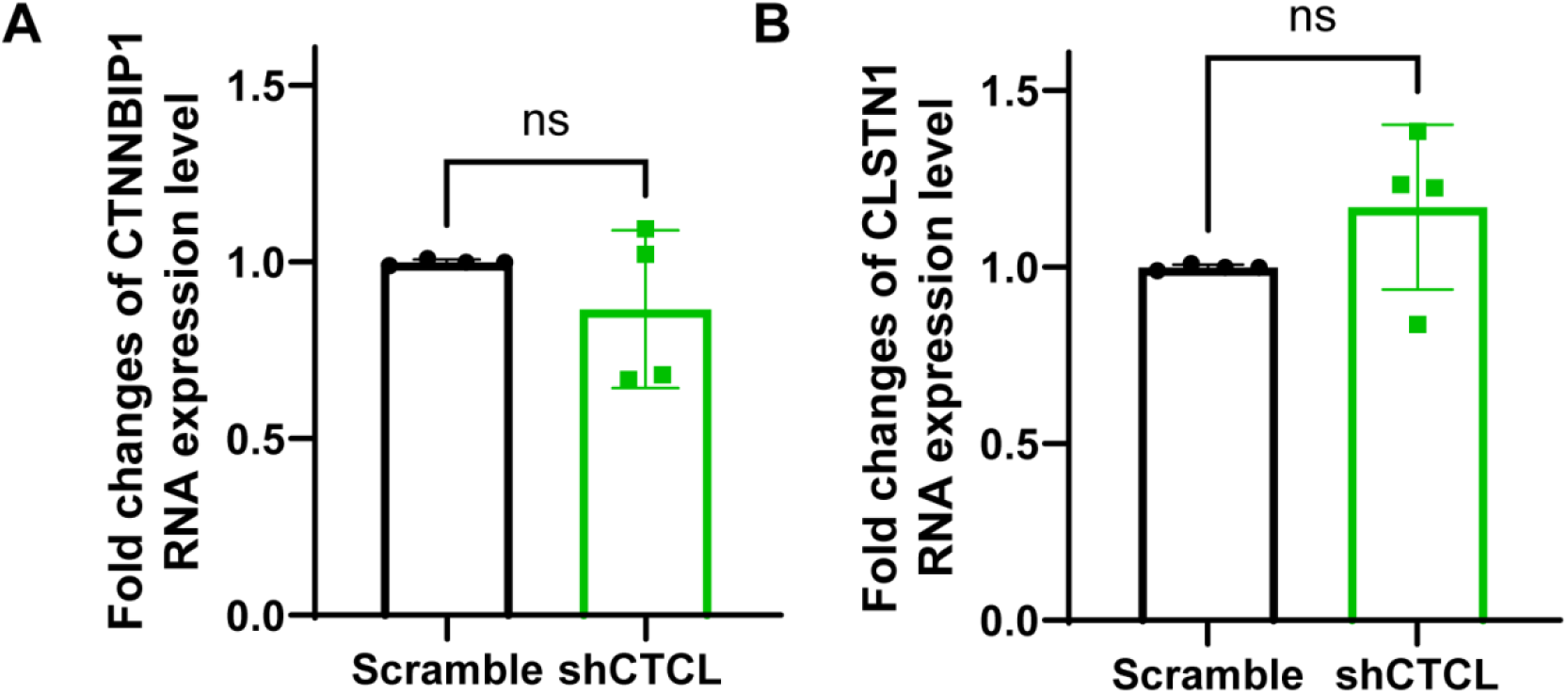
The RT-qPCR quantification of CTCL parental gene expression. (A) The knockdown experiments showed no effect on the RNA expression levels of CTNNBIP1, n = 4, *t-test, p* > 0.05. (B) The knockdown experiments showed no effect on the RNA expression levels of CLSTN1, n = 4, *t-test, p* > 0.05.

**Figure 5–figure supplement 3.**
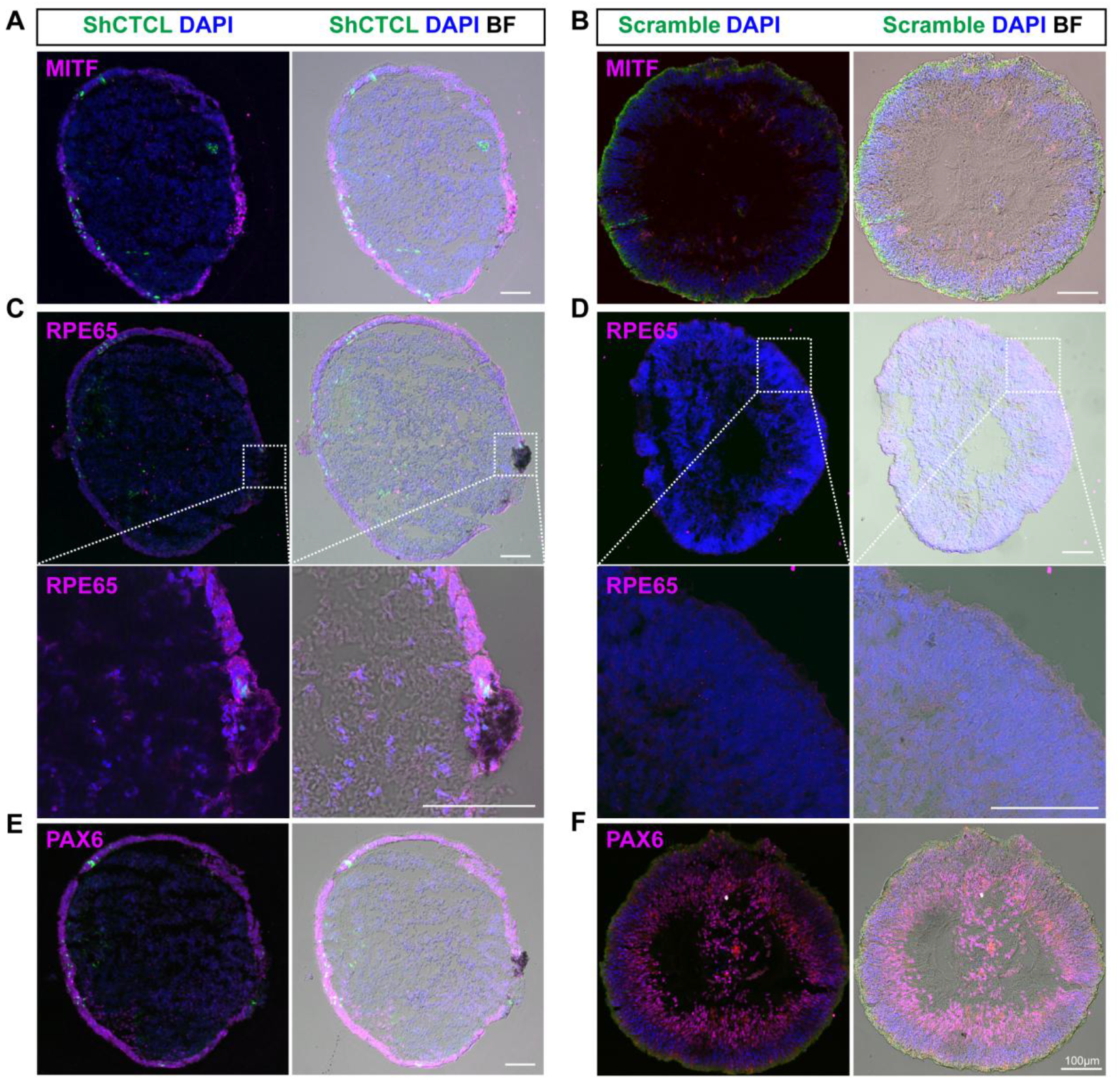
Cryosection immunofluorescence staining for D60 organoids. (A, B) Immunostaining of RPE specific marker MITF at CTCL-knockdown organoids (A) and scramble shRNA organoids (B). (C, D) Immunostaining of RPE specific marker RPE65 at CTCL-knockdown organoids (C) and scramble shRNA organoids (D). The lower panel showed the amplified images of the indicated areas. (E, F) Immunostaining of PAX6 at CTCL-knockdown organoids (E) and scramble shRNA organoids (F). Scale bars = 100μm. BF, bright-field images.

**Figure 6–figure supplement 1.**
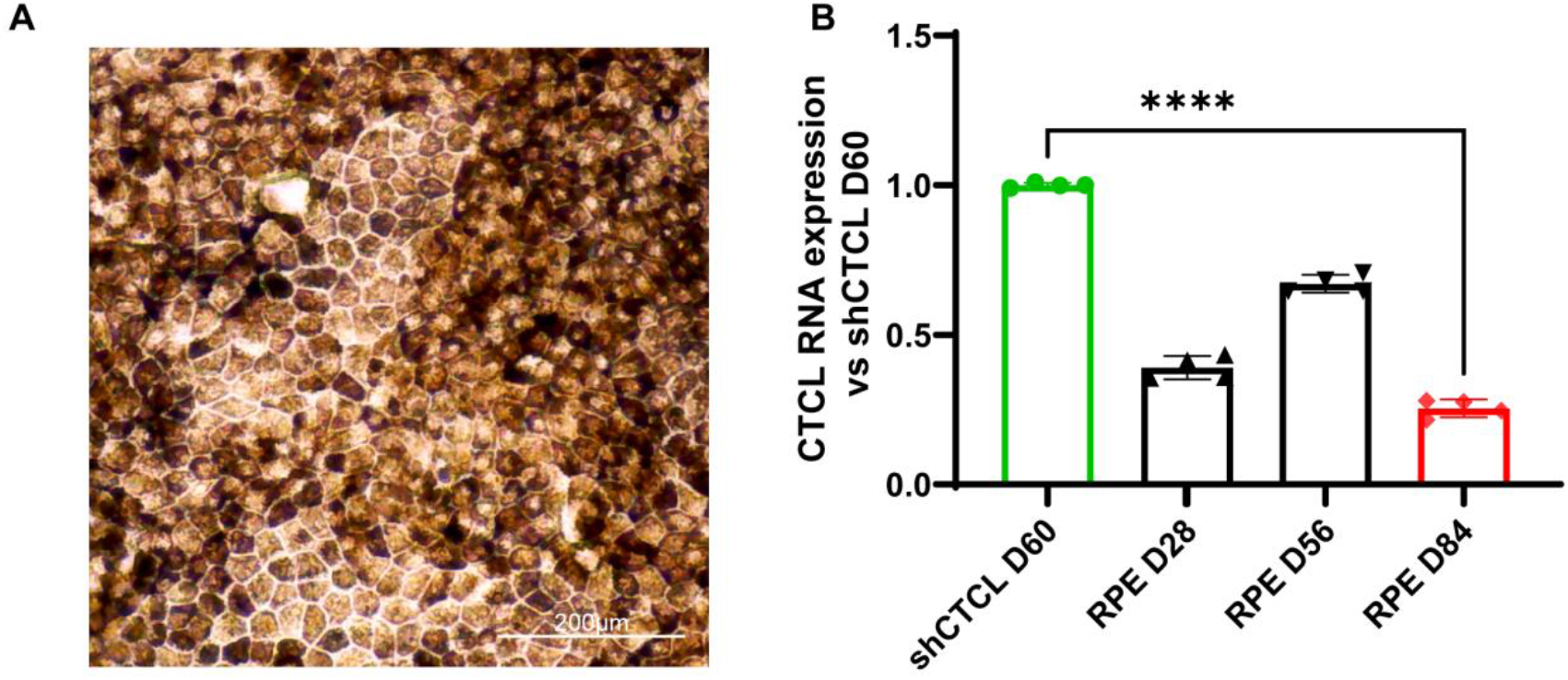
(A) Bright-filed image of mature RPE differentiated from induced pluripotent stem cells. (B) The qRT-PCR results of CTCL RNA expression levels in D60 CTCL-knockdown organoids and the differentiated RPEs at the indicated timepoints. It showed that CTCL was significantly downregulated in RPEs, even compared with the shCTCL-treated retinal organoids, n = 4, *t-test, p* < 0.0001.

